# Sexual antagonism and sex determination in three syngnathid species alongside a male pregnancy gradient

**DOI:** 10.1101/2023.08.30.555491

**Authors:** Arseny Dubin, Jamie Parker, Astrid Böhne, Olivia Roth

## Abstract

The allocation of energy towards gamete production, parental care, mate choice, sex roles, and sexual dimorphism generates divergence in selection pressures between the sexes, leading to opposing fitness strategies and sexual antagonism (SA). Due to the shared genetic makeup, a single genomic locus can contain a gene or allele with differing fitness impacts on each sex. This intralocus sexual conflict can be resolved via intersex bias in gene expression and/or formation of sex-linked genomic regions, that may also regulate sex determination. Sex determination (SD) encompasses environmental SD (ESD), monogenic SD, and polygenic SD. Occasionally, shifts from one SD locus to another can occur. While the precise mechanisms driving these shifts are unknown, SA is believed to be a major contributor. To investigate the link between SA and SD, we selected three syngnathid species along the gradient of male pregnancy that evolved with different sex roles and intensities of sexual dimorphism. By looking at intersex genetic divergence (Fst) and sex-biased expression patterns, we uncovered that sex role and mate competition, rather than male pregnancy, primarily drive SA. Furthermore, we identified processes related to non-coding RNAs and biased allele expression as mediators of SA. Most notably, we discovered intraspecies sex chromosome polymorphism in *Hippocampus erectus*. Overall, we report important details on the interplay between SA and SD, and suggest that understanding SA and its resolution mechanisms is crucial for unraveling the evolution of SD in diverse species.

## 1. Introduction

A phenotype that directly or indirectly enhances reproductive success (i.e., fitness) is expected to increase in frequency within a population through natural selection (Gregory 2009). The selection acting on a phenotype arises from a dynamic interaction between an organism and its environment. The environment influences an organism’s gene expression and embryogenesis, thereby shaping its development (Hamdoun and Epel 2007; Bosch et al. 2014; Moczek 2015). Subsequently, this affects its reproduction and survival. In turn, organisms actively interact with their surroundings, selecting and modifying their environment, which adjusts their probability of survival and reproduction (Laland et al. 1999; Trappes et al. 2022). This makes an organism both subject and object of its own evolution (Levins and Lewontin 1985; Svensson 2018; Erik I Svensson 2023). Such bi-directional cause-and-effect dynamics extend beyond the organism and its abiotic environment, and include interactions between the sexes of the same species — where one sex selects for certain phenotypes of the other sex and *vice versa* (Svensson 2018; Svensson 2019; Erik I Svensson 2023; Erik I. Svensson 2023).

The emergence of sexual reproduction fostered the formation of male and female gametes in most metazoans, resulting in a division of sexes (Bachtrog et al. 2014; Furman et al. 2020). The allocation of energy into gamete production, parental care, mate choice, sex roles and sexual dimorphism all contribute to differences in natural and sexual selection between the sexes (Parsch and Ellegren 2013; Bachtrog et al. 2014; Mank 2017; Rowe et al. 2018; Svensson 2019; Furman et al. 2020; Erik I. Svensson 2023). This variation in selection can lead to opposing male and female fitness strategies, where the optimal strategy for one sex negatively impacts the other, resulting in sexual antagonism (SA) (Rice and Holland 1997; Lucotte et al. 2016; Svensson 2018; Iversen et al. 2019; Svensson 2019; Tosto et al. 2023). This conflict transforms the relationship between the sexes into a perpetual “tug of war” – a reciprocal cause- and-effect dynamic fueled by SA. Due to the shared genome sequence between males and females, a single genomic locus can harbor a gene or allele with divergent fitness effects on the sexes. This divergence in fitness effects of a gene/allele is termed intralocus sexual conflict, and loci that harbor these genes/alleles are defined as sexually antagonistic loci (Bachtrog et al. 2014; Mank 2017; Smith et al. 2023).

Intralocus sexual conflict can be resolved via two primary mechanisms: intersex bias in gene expression or the confinement of conflicting alleles to the benefiting sex (Bachtrog et al. 2014; Cheng and Kirkpatrick 2016; Wright et al. 2018; Furman et al. 2020; Lichilín et al. 2021; Smith et al. 2023; Tosto et al. 2023). This confinement can be achieved through reduced recombination rates and the formation of sex-linked genomic regions (Bachtrog et al. 2014; Furman et al. 2020; Smith et al. 2023). While sex-biased gene expression can alleviate intralocus sexual conflict, it seems that sometimes it is not enough for its complete resolution. Genes exhibiting sex-biased expression were found to display genetic differentiation between the sexes, measured by elevated intersex Fst values (Cheng and Kirkpatrick 2016; Wright et al. 2018). Therefore, the resolution of intralocus sexual conflict appears to depend on a balance of differential gene expression and allele fixation (Bachtrog et al. 2014; Cheng and Kirkpatrick 2016; Wright et al. 2018; Furman et al. 2020; Smith et al. 2023; Tosto et al. 2023). This association links intralocus sexual conflict resolution and the evolution of sex determination (SD) systems, which also often involve formation of sex-linked genomic regions with suppressed recombination.

The mechanisms underlying SD exhibit remarkable diversity across clades and even closely related species (Bachtrog et al. 2014; Furman et al. 2020). Mammals and birds rely on generally well-conserved genetic SD system involving XY and ZW sex chromosomes, respectively (Bachtrog et al. 2014; Furman et al. 2020). In contrast, reptiles and teleost fish have evolved a broader range of SD strategies, ranging from environmental SD (ESD), to monogenic SD with a single master SD gene, and polygenic SD (Judith E Mank et al. 2006; Judith E. Mank et al. 2006; Bachtrog et al. 2014; Furman et al. 2020). The latter is believed to represent a transitional state that occurs during the shift from one SD locus to another, often referred to as a “turnover” event (Schartl et al. 2023). The observed variation in SD loci among closely related species indicates a turnover event in their evolutionary history. The frequency of turnover events varies across taxa, with some, like the teleost clade *Cichlidae*, experiencing them more frequently than others (El Taher et al. 2021). Interestingly, novel SD loci, despite evolving independently, are often co-opted from a limited subset of genes involved in the classical vertebrate sex differentiation network. Seemingly an exception to this pattern is the *de novo* evolved salmonid master SD gene (*sdY*), which arose from a duplication of IFN regulatory factor 9 (*irf9*) (Bertho et al. 2018), a gene that is not involved in sex differentiation. The *sdY* gene product was found to interact with the female-determining transcription factor *foxl2*, which belongs to the classical sex differentiation network. This interaction stops the female differentiation pathway, facilitating testis development. This confirms the notion that selection for novel SD loci in a turnover event is restricted to genes within the classical sex differentiation cascade or genes that interact with it (Sander Van Doorn 2014; Bertho et al. 2018). While the precise mechanisms driving turnover events remain unclear, SA is thought to be a major contributor. Currently, one of the main research gaps revolves around the question of precedence: what comes first, an SD locus or a region under SA? It is unclear whether an SD gene emerges first, leading to a region of recombination suppression and subsequent accumulation of SA loci within the region, or whether SA loci promote the emergence of an SD locus by selecting for a region of recombination suppression (Sander Van Doorn 2014; Schartl et al. 2023; Smith et al. 2023). SD research has proposed several models describing the relationships between SD and SA loci and the roles these relationships play in turnover events, each with varying degrees of support (Schartl et al. 2023; Smith et al. 2023).

The teleost family Syngnathidae is famous for their unconventional morphology and unique form of parental care – male pregnancy. The degree of parental investment evolved on a gradient within the family, ranging from simple egg attachment to a sealed brood pouch with a placenta-like structure to supply oxygen, and potentially nutrients, to the developing embryos (Wilson 2001; Skalkos et al. 2020). In addition to these differences in parental care among syngnathid males, there is also variation in sex roles across the species. Some species, e.g., most seahorses, adhere to “conventional” sex roles in which the females choose their mate. In other species, e.g., most pipefishes, females compete for access to mates and carry costly secondary sexual signals, while the male chooses the mate and cares for the offspring (Wilson et al. 2003). Both, varying sex roles with sexual dimorphism as well as differences in parental investment are expected to contribute to strong divergence in selection between the sexes, which in turn should be reflected in the underlying genomic and transcriptomic profile.

To investigate the link between SA and SD, we selected three syngnathid species along the gradient of male pregnancy that evolved with different sex roles and intensities of sexual dimorphism. The species with the most advanced form of male pregnancy *Hippocampus erectus* (sealed brood pouch and a placenta-like structure to nurture the eggs) has evolved with conventional sex role, monogamy, and larger males (Foster and Vincent 2004). *Syngnathus typhle* and *Nerophis ophidion* represent species with reversed sex roles and polygamy. *N. ophidion* has a strong sexual dimorphism with larger decorated females (Wilson et al. 2003). In *S. typhle*, in contrast, males and females are difficult to differentiate when out of their mating season. *S. typhle* has a more advanced form of male pregnancy with eggs protected by an inverted brood pouch, whereas *N. ophidion* males have external egg attachment on the ventral side (Wilson et al. 2003). Given the profound influence of male pregnancy on morphology, physiology, and life history, we hypothesize that male pregnancy is the primary driver of SA relationships, and has a profound effect on the evolution of SD systems in these species. We anticipate either a strong genetic sex-linkage or a pronounced divergence in expression patterns of genes associated with adaptation to male pregnancy, e.g., those involved in osmotic regulation, oxygen transport, and immune function. We expected the extent of genetic and expression divergence to correlate with the gradient of male pregnancy complexity: *N. ophidion* with the weakest SA signal, *S. typhle* with intermediate signal and potentially a sex-linked region, and *H. erectus* with the strongest signal and a large sex-linked genomic region. Alternatively, a different pattern would suggest that other factors, such as dimorphism and sex roles, also play significant roles. To look for signatures of SA, we used RNA-seq data obtained from head kidney and gills of males and females of each species. In addition, we investigated the relationship between SA loci and an SD system by searching for distinct regions of elevated genetic divergence between the sexes (intersex Fst) and the accumulation of genes/alleles showing sex-bias in expression in these regions.

This study sheds light on the underlying processes involved in SA in three syngnathid species, sex-linked genomic regions in these species, and the relationships between the loci under SA and sex-linked regions.

## 2. Methods

### 2.1 Sample collection, nucleic acid isolation and sequencing

An adult *Syngnathus typhle* male and an adult *Nerophis ophidion* female, were taken from our animal facility at GEOMAR, Germany. Large animals were chosen to achieve sufficient amounts of high molecular weight (HMW) DNA for Pacific Biosciences library preparation. *N. ophidion* displays a strong sexual size dimorphism with large females and small males. We thus sampled a female *N. ophidion* for genome sequencing.

DNA extraction, library preparation, and PacBIO sequencing was performed by the Norwegian Sequencing Centre, Oslo, Norway (sequencing.uio.no).

The DNA samples were processed according to the Pacific Biosciences Express library preparation protocol without prior fragmentation. The final libraries were size-selected with a BluePippin with 11 kb cut-off and sequenced on a Sequel II instrument using a Sequel II Binding kit v2.0 with Sequencing chemistry v2.0. Circular Consensus Sequences (CCS) were generated using the CCS pipeline (SMRT Link v10.1.0.119588) with default settings.

*Hippocampus erectus* individuals were bred at the GEOMAR aquaria facilities, Kiel, for several generations prior to the experiment. *Syngnathus typhle* and *Nerophis ophidion* were wild-caught in the Baltic Sea (54°23′N; 10°11′E, Germany) and subsequently kept in aquaria facilities at GEOMAR Helmholtz Centre for Ocean Research, Kiel, Germany. Ten males of each species were used. Five females were used in the case of *N. ophidion* and *S. typhle*, and seven in the case of *H. erectus*. All fishes were euthanized with an overdose of MS-222 (Tricaine, 500 mg L−1; Sigma-Aldrich). After dissection, head kidney and gill samples were collected from each individual and placed in an RNAlater solution (Invitrogen) and incubated at 4°C for one week. Samples were then stored at −20°C until extraction. Total RNA was extracted using an RNeasy Mini Kit (Qiagen) according to the manufacturer’s protocol. Libraries were prepared using Illumina TruSeq stranded mRNA kit (2×150bp) and sequenced on an Illumina NovaSeq 6000 instrument at the Competence Centre for Genomic Analysis (CCGA), Kiel.

### 2.2 Genome and transcriptome assembly and annotation

#### 2.2.1 Genome assembly

The *S. typhle* and *N. ophidion* HiFi reads (CCS reads with >99% accuracy) were assembled using the Genome Assembly pipeline (SMRT Link 10.1.0.119588) with default settings. Gene space completeness of the assemblies was assessed with BUSCO (v5.2.2) (Manni et al. 2021) using the Actinopterygii reference protein set. Continuity of the assemblies was assessed with Quast (v5.0.2) (Mikheenko et al. 2018). The chromosome-level assembly of *H. erectus* and supporting annotation used in this study was previously published by Li et.al (PRJNA613176) (Li et al. 2021).

#### 2.2.2 Transcriptome assembly and gene prediction

The RNA reads generated in this study were quality trimmed with Fastp (v0.20.1) (Chen et al. 2018) with default settings. Reads from both organs were pooled and assembled with Trinity (v2.8.1) (Grabherr et al. 2011), to generate a single transcriptome assembly per species. In order to reduce redundancy of transcripts per gene, the resulting assemblies were subjected to several filtering steps. First, assemblies were filtered with Transrate (v1.0.3) (Smith-Unna et al. 2016) using raw reads as evidence. The software assigns a quality score for the entire assembly and per contig on the basis of various metrics, like the base quality scores, number of reads mapped per contig etc., determines an optimal threshold and filters the assembly. Transrate-filtered assemblies were subjected to CD-Hit (v4.8.1) (Li and Godzik 2006) clustering transcripts with 98% similarity threshold into one. These contigs were then used to identify putative protein sequences encoded within the assemblies with TransDecoder (v5.5.0). Obtained protein sequences were clustered with CD-Hit (v4.8.1) (Li and Godzik 2006) with 95% similarity threshold. Sequence identifiers of these filtered proteins were used to filter the Transdecoder CDS fasta file to reduce redundancy by only selecting the CDS sequences whose protein product passed a similarity filter. We used these filtered cDNA sequences as our transcriptome assemblies in all subsequent analyses.

#### 2.2.3 Repeat masking and genome annotation

Prior to genome annotation, repetitive elements were identified and masked. EDTA (v2.0) (Xu and Wang 2007; Gremme et al. 2013; Xiong et al. 2014; Ou and Jiang 2018; Ou et al. 2019; Shi and Liang 2019; Su et al. 2019; Zhang et al. 2022) was used to produce a *de novo* repeat library for *S. typhle* and *N. ophidion* assemblies. The resulting repeat libraries were used to mask the corresponding assemblies with RepeatMasker (v4.1.2-p1) using the settings “-a -gff - pa 32 -u -xsmall -nolow”.

To use RNA-sequencing data as evidence for gene prediction, for both *S. typhle* and *N. ophidion,* RNA reads were aligned to their respective species’ genome assemblies with STAR (v2.7.9a) (Dobin et al. 2013) in two-pass mode. Only uniquely mapped reads were considered. Filtered transcriptome assemblies were aligned to the genomes with GMAP (version 2017-11-15) (Wu and Watanabe 2005) with the following options “-B 5 -t 32 --input-buffer-size=1000000 --output-buffer-size=1000000 -f samse”.

The *S. typhle* and *N. ophidion* genome assemblies were annotated using BRAKER (v2.1.6) (Lomsadze 2005; Stanke et al. 2006; Gotoh 2008; Stanke et al. 2008; Li et al. 2009; Barnett et al. 2011; Iwata and Gotoh 2012; Lomsadze et al. 2014; Buchfink et al. 2015; Hoff et al. 2016; Hoff et al. 2019; Brůna et al. 2020; Brůna et al. 2021). We ran BRAKER twice. First with RNA, both the unassembled reads and the transcriptome assemblies, then with OrthoDB proteins (odb10_vertebrata_fasta) (Kriventseva et al. 2019) as evidence. Afterwards, we combined the annotations with TSEBRA (v1.0.3) (Gabriel et al. 2021), giving the RNA evidence more weight (pref_braker1.cfg). The cDNA and amino acid sequences were obtained from the final BRAKER GTF with AGAT (v0.8.0) (Dainat et al. 2023) toolkit using agat_sp_extract_sequences.pl script with “--aa --clean_final_stop –clean_internal_stop” options to get amino acid sequences, and “-t exon –merge” to get cDNA. The putative function of predicted protein sequences was determined with Orthofinder (v2.4.0) (Emms and Kelly 2019) using sequences from the following fish species: *Danio rerio (GRCz11), Gasterosteus aculeatus (BROADS1), Hippocampus comes (H_comes_QL1_v1, GCA_001891065.1), Hippocampus erectus (PRJNA613176), Lepisosteus oculatus (LepOcu1), Oryzias latipes (ASM223467v1), Syngnathus acus (GCF_901709675.1_fSynAcu1.2), Takifugu rubripes (rubripes.fTakRub1.2), Syngnathus rostellatus* (GCF_901709675.1).

#### 2.2.4 Transcriptome functional annotation

The transcriptomes were functionally annotated in a similar manner to the genomes. We used predicted proteins from *H. erectus*, *S. typhle* and *N. ophidion* transcriptome assemblies together with the previously mentioned eight fish species to run orthology analysis with Orthofinder (v2.4.0) (Emms and Kelly 2019).

### 2.3 Variant calling

#### 2.3.1 Genome-independent approach

A similar approach as described in El Taher et al. (El Taher et al. 2021), KisSplice (v2.4.0) (Sacomoto et al. 2012; Lopez-Maestre et al. 2016; Benoit-Pilven et al. 2018) was used to call SNPs directly from raw RNA reads without prior genome mapping. The SNP data was then processed with the kissDE R package (v1.14.0) (Sacomoto et al. 2012; Lopez-Maestre et al. 2016; Benoit-Pilven et al. 2018) to identify SNPs that showed differential expression between sexes. To predict their functional impact and identify the genes that exhibited differential allele expression, the SNPs were placed on the corresponding transcriptome assemblies with BLAT (v36) (Kent 2002) using “–minIdentity=80” option and KisSplice2refTranscriptome (v1.3.3) (Sacomoto et al. 2012; Lopez-Maestre et al. 2016; Benoit-Pilven et al. 2018). At this step, we utilized KisSplice2refTranscriptome flags to filter the data, which excludes SNPs that occurred in multiple assembled genes (SNP_in_mutliple_assembled_genes flag) and could simply represent a sequencing error (Possible_sequencing_error flag). Then, we filtered the dataset to include only bi-allelic SNPs. Moreover, we applied filters to include SNPs that occurred only in CDS.

As we wanted to investigate both, sex-biased allele expression and identify sex-linked regions within the genome, the aforementioned SNPs were divided into two categories: i) differentially-expressed alleles (DEA) – alleles that show a bias in expression, i.e., differences in read counts for pairs of variants, between the sexes. ii) Sex-specific alleles (SSA) (male or female) – were defined as SNPs where all individuals of one sex are homozygous at this locus, whereas at least 75% of individuals of the opposite sex are heterozygous at this locus. We considered these as fixed differences but point out that due to the usage of RNA sequencing data those could also represent extremes of differential allele expression. However, the KisSplice developers previously showed that expressed allele frequencies very closely reflect true allele frequency differences as would be derived from DNA sequencing (Sacomoto et al. 2012; Lopez-Maestre et al. 2016). In the following, with SNPs of category ii) we considered two potential sex determination systems – an XY system with a male-specific Y and ZW system with a female-specific W chromosome.

#### 2.3.2 Genome-based approach

The RNA reads were aligned to the annotated genomes with STAR (v2.7.9a) (Dobin et al. 2013) in two-pass mode with the options “--outFilterIntronMotifs RemoveNoncanonical -- outSAMunmapped None --outFilterMultimapNmax 1”. The alignment data was then processed with the Genome Analysis Toolkit (GATK) (v4.2.2.0-17) (Poplin et al. 2017) and Picard (v2.25.4) (Broad Institute 2019). First, duplicate reads were marked (MarkDuplicates) and reads with Ns in the cigar strings were split into multiple alignments (SplitNCigarReads). Then, potential variants were called on each sample in gVCF mode (–ERC GVCF option). gVCF files for each sample were combined into a single gVCF per species and the variants were called with GATK’s joint genotyping method.

As GATK guidelines do not recommend to use standard hard filtering criteria when working with SNPs called from RNA-seq data, the final VCFs were filtered with “QD < 2.0” and “FS > 30.0” thresholds applied, and using “--select-type-to-include SNP --restrict-alleles-to BIALLELIC” options to keep only high quality bi-allelic SNPs.

#### 2.3.3 Fst calculation

Using SNPs resulting from the genome-based approach, we calculated intersex Weir and Cockerham Fst values for each of the genomes in windows of three sizes – 1Kb, 10Kb, 100Kb with VCFtools (v0.1.14) (Danecek et al. 2011). To minimize bias introduced by uneven numbers of males and females, we used data from only 5 individuals of each sex and set a missing data threshold to 20 percent.

#### 2.3.4 Overlap between approaches

In order to minimize software-related bias, we investigated the overlap between SNPs called by KisSplice and GATK. To place KisSplice-called variants on a genome, we aligned KisSplice SNPs to the genome using STAR (v2.7.9a) (Dobin et al. 2013) and processed these alignments with KisSplice2refgenome (v2.0.5) (Sacomoto et al. 2012; Lopez-Maestre et al. 2016; Benoit-Pilven et al. 2018). Alleles and their locations were compared using a custom python script.

#### 2.3.5 Differential gene expression analysis

Transcript abundance was calculated with Kallisto (v0.46.1) (Bray et al. 2016) and imported into DESeq2 (v1.22.2) (Love et al. 2014) with tximport (v1.22.0) (Soneson et al. 2015). Genes with total counts less than 10 across all samples were removed from the dataset. Expression between sexes was analyzed using DESeq2 per species and organ with sex representing the model contrast. The alpha value was set to 0.01. The difference in expression was considered significant if an adjusted p-value was lower than 0.01. The resulting gene lists were used in conjunction with Fst scans to identify genes that are differentially expressed and located within a region of elevated Fst.

#### 2.3.6 Intersex SNP density analysis – kmerGWAS analysis

For both, SNP density calculations and kmerGWAS analysis, we followed the SexFindR workflow suggested by Grayson and colleagues (Grayson et al. 2022).

The SNP density per sample was calculated with vcftools in 10kb windows on bi-allelic SNPs called by GATK from the previous steps. The results were analyzed using SNPdensity_permutations_fugu.R script, provided within the SexFindR workflow.

The script calculates the mean number of SNPs in males and females per window and then calculates the difference between the male and female means.

The KmerGWAS analysis was performed according to the SexFindR recommendations with no modifications, using kmersGWAS library (Voichek and Weigel 2020), PLINK (v1.07) (Purcell et al. 2007), ABySS (v2.0.2.) (Jackman et al. 2017), and BLAST (v2.5.0) (Altschul et al. 1990; Camacho et al. 2009; Boratyn et al. 2012). Results were visualized with R.

## 3. Results

### 3.1 Genome assemblies and annotation

To identify sex-linked genomic regions and provide additional support for genome-independent SNP calling, we generated draft genome assemblies and their annotations for two of the studied species. While the here used *H. erectus* genome was already published (Li et al. 2021), the *N. ophidion* and *S. typhle* genomes were newly generated for this study. From here, the species will be referred to as HE (*Hippocampus erectus*), NO (*Nerophis ophidion*), and ST (*Syngnathus typhle*).

The NO genome was sequenced to ∼22x coverage with PacBio HiFi reads of 14.9 Kb mean read length. The resulting assembly length was 1.7 Gb with an N50 of 1.2 Mb. For the ST genome, the coverage with PacBio HiFi was ∼40x with a mean read length of 14.7 Kb. The total length of the assembly was 405 Mb with an N50 of 2.3 Mb. While we did not reach chromosomal resolution with these assemblies, the N50 of 1.2 and 2.3 Mb allowed us to detect large sex-linked regions, blocks of genes with sex-biased allele expression patterns and their synteny (i.e., corresponding linked locations).

The BUSCO gene space completeness scores revealed high levels of completeness with the *Actinopterygii* dataset. HE: Complete:96.6%/Missing:2.3%, NO: Complete:93.9%/Missing:4.0%, and ST: Complete:94.4%/Missing:4.1% respectively. The BRAKER2 workflow resulted in 27’499 genes annotated in ST (BUSCO C:93.7%/M:4.8%), and 38’598 genes in NO (BUSCO C:93.2%/M:4.3%). For comparison, the current HE annotation contains 20’137 genes (BUSCO C:97.4%/M:1.7%). The inflated gene number in NO could be attributed to the extremely high repeat content of the genome which makes gene prediction difficult even when the genome is masked (Roth et al. 2020), resulting in erroneous predictions. This inflated gene number bias was to some extent corrected by the orthology assignment. 19’762 genes (98.1%) of HE were attached to orthogroups, 29’875 (77.44%) of NO, and 23’907 (86.9%) of ST. To facilitate comparisons across species and their genomes, in the subsequent analyses we only used genes that had orthologs to the zebrafish *Danio rerio –* 21’685 in HE, 21’801 in NO, and 20’362 in ST. This reduced the bias in gene numbers across our species set.

### 3.2 Alignment of variant calling approaches

We used SNP data to assess allele frequency differences between the sexes and identify putative sex-linked regions. For each part, however, a different variant calling approach was applied. We wanted to avoid that assembly and annotation quality influence differential allele expression results and thus opted for a genome-independent variant calling approach in this part of the study. GATK-derived reference genome-based data was used to search for sex-linked regions within the genome and provided additional support for SNPs called with the genome-independent approach. To compare the two approaches, we placed pre-filtered bi-allelic SNPs called with KisSplice on the genomes. With a custom python script, we compared both, location of the SNP and the called allele. We found that around 79 to 83 % of KisSplice SNPs were confirmed by the GATK bi-allelic SNPs suggesting high congruence.

### 3.3 Differential allele expression and sex-linked SNPs

#### 3.3.1 Differential allele expression between the sexes

We here considered a scenario where an intralocus sexual conflict would cause a difference in allele expression. Sex differences in allelic expression point towards an emerging or ongoing sexual conflict. The accumulation of sex-specific alleles (SSAs) at certain locations in the genome supports actual sex-linkage of the underlying SNPs/alleles linked to the evolution of a sex chromosome. Sex chromosomes can ultimately resolve sexual conflicts by confining SA genes/alleles to the sex they benefit. Species with XY sex determination systems are expected to have considerably more or only male-specific SNPs due to their male Y chromosome, while ZW species are expected to have more or only female-specific SNPs.

Here, allele level differential expression between the sexes was looked at specifically. We categorized bi-allelic SNPs into SSAs male or female, and differentially expressed alleles (DEAs).

All three examined syngnathid species showed signs of sex-specific allele expression. The number of DEAs was considerably higher in NO than in the two other species assessed (Ratio: 3:1) (Supplementary tables S4,S6,S9). Next, we looked at species-specific distributions of male and female SSAs. While male and female SSAs were identified in NO and ST, HE only harbored male SSAs (Figure 1. Panels A in each species, Supplementary tables S5, S7, S8,S10,S11). Overall, in our analysis, ST exhibits the weakest divergence between the sexes in both DEA and SSA numbers. On the other side of the spectrum, HE had ∼100 times more male SSAs than ST and ∼17 times more than NO. To examine the genomic locations of these male SSAs identified using transcriptome assemblies, genome and transcriptome annotations were matched. The HE *chr14* appeared to harbor ∼2.4 times more genes with male-specific alleles than the next best hit (124 vs. 52). With the exception of *chr14*, other notable hits came from unplaced scaffolds. Upon further examination, it appeared that both the *chr14* and the aforementioned scaffolds share all their 1–1 orthologs with *chr1* of *Hippocampus abdominalis* (HA) suggesting that they all belong to the same chromosome. With that in mind, the expanded *chr14* would be the size of ∼34.3Mb (vs 13.1Mb original length), which is similar to ∼36 Mb of its supposed homologue – the *chr1* of HA.

**Figure 1.**
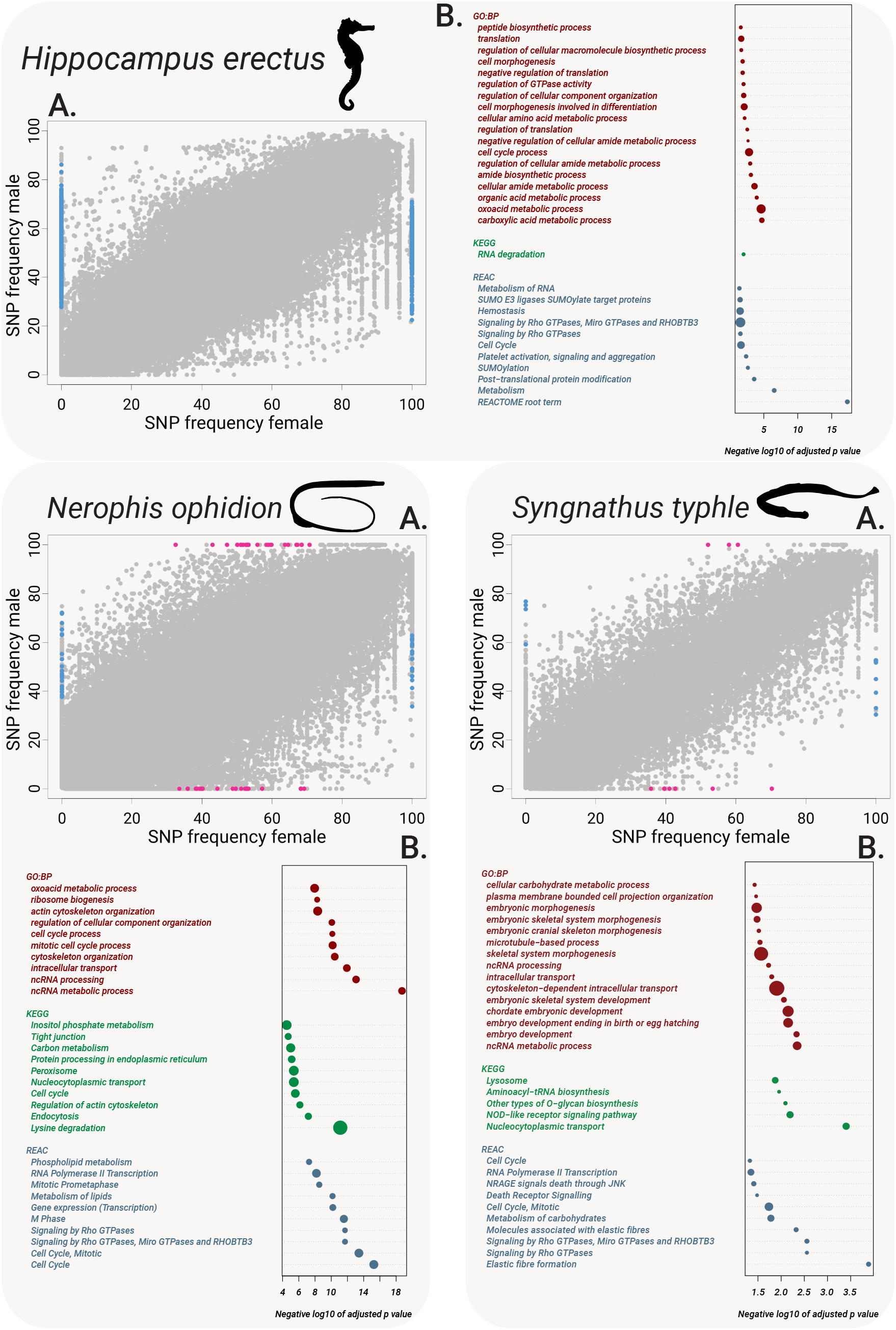
Panels - *Hippocampus erectus*, *Nerophis ophidion*, *Syngnathus typhle*. **Subpanel A** - male vs female allele frequencies of Kissplice-called SNPs. Coloured grey are the variants that show neither sex-specificity, nor differential expression between sexes. Marked with blue are male-specific SNPs, marked with pink - female specific SNPs. **Subpanel B** - results of an enrichment analysis for 3 data sources: GO BP (red), KEGG (green) and Reactome (blue). Each point represents an enriched term, with name of the term located on the Y-axis, and padj values on the X-axis. The size of the point represents a relationship between the number of genes from the list that belong to this term vs the term size. Meaning, point size represents a proportion of the term is that is represented but the overlap genes.

The expanded *chr14* would contain 327 genes that bear male-specific alleles (∼9.5 per Mb) compared to the now next best 15 hits from *chr4* (∼0.7 per Mb). This pattern marks HE as a candidate XY species.

To assess biological processes and pathways that might be affected by differential allele expression, we applied an over-representation analysis (ORA) to gene lists that contained DEAs (Figure 1. Panels B, for each species). The analysis suggested “Cell Cycle”, “ncRNA metabolic process”, and “Signaling by Rho GTPases” to be overrepresented in gene lists with DEA in all three species. The broad KEGG term “Cell Cycle” includes 469 genes involved in cell replication and nuclear division. The pathways involved in RNA processing and metabolism also appeared overrepresented in all species. The ncRNA metabolic pathway is affected by differential allele expression in all studied species, and ncRNA processing only in NO and ST. Some more broad GO-terms “RNA degradation”, “Metabolism of RNA”, “tRNA metabolic process”, however, appeared only in some species. The third overrepresented pathway – “Signaling by Rho GTPases” is involved in many cellular processes and helps cells to respond to external and internal stimuli via Rho GTPases – a family of molecular-switch proteins (G proteins) (Bar-Sagi and Hall 2000; Jaffe and Hall 2005; Hodge and Ridley 2016; Perry and Maddox 2019; Boueid et al. 2020). Several members of the G proteins family were found to be crucial for spermatogenesis (as examined in mammals) (Lui et al. 2003; Zhang et al. 2010; Zhu et al. 2023). Processes related to noncoding RNAs (ncRNA metabolic process and ncRNA processing) were found to be influenced by DEA between sexes in all species.

HE and NO share a higher number of pathways affected by sex-biased allele expression – mainly cell maintenance, signal transduction, intracellular transport, and various metabolic processes (Figure 1. Panels B, for mentioned species). While ST had some overrepresented pathways involved in cell cycle and metabolism, the main affected pathways appeared to be related to embryonic morphogenesis and embryo development. Such a reproduction-related pathway overrepresentation among DEAs might be particularly relevant to understanding sexual conflict in ST, considering that ST exhibits the weakest differences between sexes in both DEA and SSA numbers. It is difficult to judge, however, what roles these pathways play in the sampled organs.

The NOD-like receptor signaling pathway is enriched in genes with DEAs in NO and ST. Among them were NLR family member X1 (*nlrx1*), nucleotide-binding oligomerization domain-containing protein 2 (*nod2*) and TNF receptor associated factor 2b (*traf2b*) in NO, TNF receptor-associated factor 5 (*traf5*) and interleukin-1 beta (*il1b*) in ST. The nucleotide-binding oligomerization domain-like (NOD-like) receptors are a type of pattern recognition receptors that play a key role in innate immune system responses (Jones et al. 2016; Chou et al. 2023).

#### 3.3.2 Sex-Specific Alleles (SSAs)

The only species that had enough genes with SSAs to conduct a meaningful enrichment analysis was HE. From a list of 353 genes with male SSAs (Supplementary tables S5), 5 pathways appeared to be overrepresented; from GO:BP – glycerol-3-phosphate metabolic process, from KEGG – VEGF signaling pathway, C-type lectin receptor signaling pathway, Herpes simplex virus 1 infection, and from Reactome – Adaptive Immune System.

Genes from “glycerol-3-phosphate metabolic process” were two glycerol-3-phosphate dehydrogenases and glycerol kinase 3. Most of the genes in overrepresented adaptive and innate immune pathways were related to signal transduction, metabolism and cell cycle, and only indirectly to the immune system. However, some genes with SSAs could support the suggested sexual immune dimorphism (Roth et al. 2011; Pappert et al. 2023), namely: integrin alpha 5 (*itga5*), caspase-9 (*casp9*), mechanistic target of rapamycin kinase (*mtor*), nuclear factor of activated T cells 2a (*nfatc2a*), and сluster of differentiation 40 (*cd40*).

Both ST and NO had genes with male and female SSAs. In NO – 24M/24F genes (Supplementary tables S7-8), and in ST 6F/4M (Supplementary tables S9-10). Since a meaningful enrichment analysis is impossible with the low number of genes involved, we scanned the lists manually and looked for genes that might be involved in SA.

Considering sampled tissues functions, we expected to see signatures of SA in genes related to the innate and adaptive immune systems in both tissues, and cellular transport in gill, since fathers might need more oxygen when they are pregnant (Goncalves et al. 2015). Notable hits included: huntingtin (*htt*, NO-F), cryptochrome circadian regulator 2 (*cry2*, NO-F), macroH2A.1 histone (*macroh2a1*, NO-F), T cell immune regulator 1, ATPase H+ transporting V0 subunit a3b (*tcirg1b*, NO-M), transporter associated with antigen processing 1 (*tap1*, NO-M), cathepsin K (*ctsk*, NO-M), ATP binding cassette subfamily B member 9 (*abcb9*, NO-M), and integrin beta 4 (*itgb4*, ST-F).

### 3.4 The search for sex-linked genome regions

We next integrated results from our sex-biased SNPs analysis above with whole-genome intersex Fst scans. Genes were sorted according to their Fst and density across the genome. We further compared our results to the study by Long et al. that focused on sex chromosomes in two seahorse species, *Hippocampus erectus* and *Hippocampus abdominalis*, and the pipefish species *Syngnathus scovelli* (Long et al. 2023). This required additional kmerGWAS analysis.

#### 3.4.1 Genome-wide picture

We calculated intersex Fst values for all three genomes in windows of 1kb, 10kb and 100kb. The 10kb and 100kb scans provided an overall look at the genome and facilitated the identification of regions or chromosomes with elevated Fst values that might be indicative of sex-linkage. If a sex chromosome or sex-linked region within a genome has accumulated sufficient fixed differences between the sexes, we would expect to see a distinct Fst peak at that location. For Weir Fst estimation we expected 0.5 Fst values for a fixed difference between the homozygous and the heterozygous sex (Gammerdinger et al. 2020). For 10kb and 100kb windows we used the threshold of 0.4. We considered a lower 0.4 threshold since values were calculated over large genomic regions that might include sites with low Fst, hence lowering the overall Fst in a given window. Each species exhibited a unique distribution pattern of Fst values across the genome, as shown in Figure 2A. ST exhibited the lowest overall Fst values, with regions of elevated Fst being sparsely distributed along the genome (33 10Kb windows with Fst >=0.4 or ∼0.08 per Mb). This finding aligns with our observations on sex-biased allele expression and complements the recently published data by Long et al. (Long et al. 2023), which focused on a different species within the same genus, *Syngnathus scovelli*. Both NO and HE had similar overall Fst levels, with NO having 1222 10Kb >=0.4 windows (∼0.7 per Mb) and HE 721 10Kb >=0.4 windows (∼1.7 per Mb).

**Figure 2.**
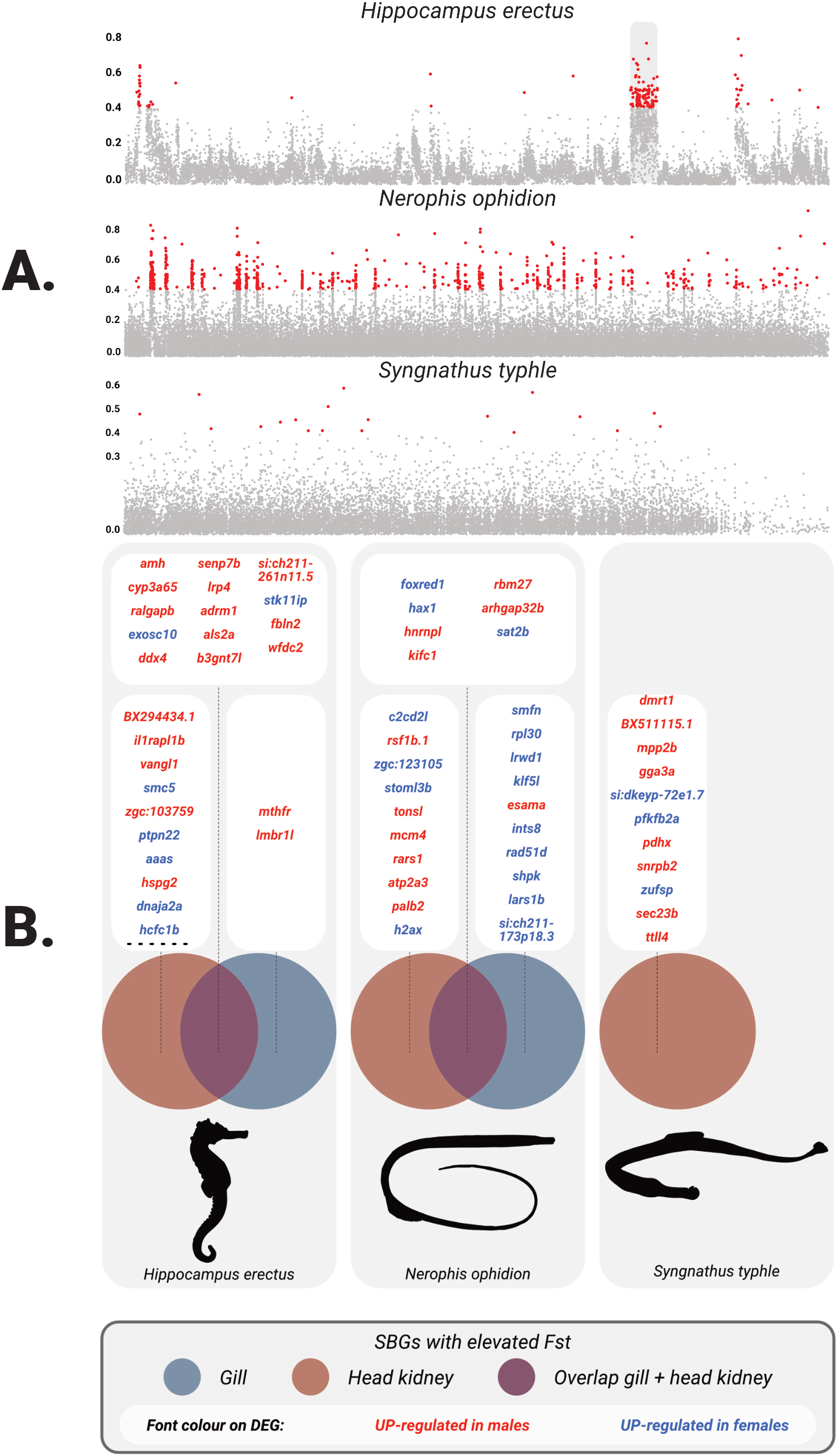
Panel A - Fst values per 10 kb windows across the genome in *H.erectus*, *N.ophidion*, *S.typhle*. Y-axis - Fst value, X-axis -position within a genome. Colored points - ones that pass the 0.4 threshold. On HE plot, grey rounded rectangle marks candidate sex chromosome. **Panel B** - the overlap between genes with elevated Fst and DEG in *H.erectus*, *N.ophidion*, *S.typhle*. The *H. erectus* gill panel includes only top 10 most significant hits indicated by the dotted line. Blue colored circles represent gill DEGs, red colored circles represent head kidney DEGs, purple - the overlap between lists of genes.Each panel with gene lists is connected to the appropriate overlap sector with a dotted line. The lists are ordered based on padj value and colored based on sex (red– up-regulation in males, blue – up-regulation in females).

However, in NO the elevated regions were scattered throughout the genome, while HE had three chromosome-size scaffolds differentiating from the rest of the genome with *chr14* having the highest number of windows with elevated Fst (183 windows or ∼13.9 per Mb) and the next two *chr1* and *chr18* with fewer hits (31 and 21 windows or ∼1.2 and ∼2.1 per Mb respectively). The fragmented nature of the genome assembly and the genome size could be responsible for sparse distribution of elevated Fst windows in NO, i.e., most of the contig hits can potentially come from a single chromosome. But considering that NO possesses both male and female SSAs (like ST), we think it is unlikely that there is a single sex-linked region or chromosome.

#### 3.4.2 Elevated Fst gene scan

The results from a 1kb-window scan were used to identify individual genes that overlap with elevated Fst windows. Using this more sensitive approach putative sex-linked genes can be identified. We used the same approach to determine the Fst threshold as with previous scans. For HE and NO we used 0.5 threshold for the reasons described above and for ST we decreased it to 0.33 to see slight bias in allele frequencies (Supplementary tables S1-3). Whenever an annotation of a particular gene overlapped with an elevated Fst window, this gene was marked and the window was annotated as belonging to this gene.

Matching the above-described scans, there was no distinct region of elevated Fst. Both NO and HE had genes with intersex Fst >0.9, which implies near-complete fixation of alleles in each sex, i.e., both sexes are homozygous at this locus and possess different alleles. Specifically, in NO, three genes demonstrated this pattern: proteolipid protein 1b (plp1b – 1.0 Fst, genomic location – ctg.000011F), tetratricopeptide repeat protein 3 (ttc3 – 0.92 Fst, genomic location – ctg.005052F) and transcobalamin beta b (tcnbb – 0.91 Fst, genomic location – ctg.000011F) (Supplementary tables S2). Similarly, in HE, near-complete allele fixation was observed for two genes: interferon regulatory factor 2 (irf2 – 0.92Fst, genomic location - chr18), and most notably anti-Müllerian hormone (amh – 0.91 Fst, genomic location – *chr13*) (Supplementary tables S1). Neither of those genes belong to the same contigs, or form a distinct cluster of genes with elevated Fst. In ST, the gene with the highest value was trafficking kinesin protein 2 binding (*trak2* – 0.8 Fst, genomic location – *ctg.000074F*) (Supplementary tables S3).

#### 3.4.3 Fst-SBGs overlap

Genes that show intersex bias in expression were also found to have elevated intersex Fst. In humans and flies the Fst scores of SBGs follow a “twin peaks” pattern, i.e., genes with an intermediate bias in expression show the highest intersex Fst, interpreted as targets of ongoing sex-specific selection on viability (Cheng and Kirkpatrick 2016).

Resolution of sexual conflict via expression bias may be insufficient in some genes, causing alleles to diverge between the sexes. SBGs with elevated intersex Fst in guppies, however, were suggested to be involved in the formation of sex-specific genetic architecture rather than intralocus sexual conflict (Wright et al. 2018).

To identify genomic signatures of SA, we investigated an overlap between SBGs and the windows with elevated Fst. Next, to assess sex-linked regions, we investigated clustering of genomic locations of SBGs with elevated intersex Fst. Because differential gene expression analysis was done using transcriptome assemblies and Fst scans using genome annotations, the results had to be compared using orthology assignment. The search was first executed per organ and then for both organs combined.

The results support the previous Fst window-scan approach. As such, in HE, the same three chromosomal scaffolds appeared to contain the most SBGs with elevated Fst: *chr14* (196 hits or ∼14.9 per Mb), *chr18* (46 hits or ∼4.6 per Mb), and *chr1* (28 hits or ∼1.15 per Mb). Among scaffolds with a high number of SBGs and elevated Fst, we identified many unplaced scaffolds, e.g., *ctg37* (128 hits), *ctg26* (92 hits), *ctg27* (67 hits), *ctg33* (54 hits), *ctg39* (19 hits). However, since our synteny analysis with the HA genome showed that all these scaffolds share 1-1 orthologs with *chr1* of HA, it is most likely that they belong to the same HE chromosome – *chr14*. The number of overlapping genes declined from here onwards. Notably, one of the well-known sex-determination genes, anti-Mullerian hormone (*amh*), appears among the most male-biased genes in both gill and head kidney of HE, and is also among the genes with the highest Fst in the genome – 0.91 (Figure 2B). The *amh* gene region, however, does not belong to either of the two candidate sex-linked scaffolds. As such, the *amh*-containing scaffold appeared during Fst scans of the genome but was among candidates with a low number of elevated Fst-windows (>300 in the top scaffolds vs 5 in the *amh*-containing scaffold).

In NO the number of SBGs with elevated Fst was not as significant as in HE, and there were more contigs that contained similar numbers of overlapping genes, which could be a consequence of a fragmented genome. There were no known sex-determining genes in the overlap.

ST showed the smallest number of SBGs with elevated Fst, and they could only be identified in the differential gene expression analysis from the head kidney. As in other species, we could not narrow it down to a single distinct region or contig enriched in overlapping genes. Most of the overlapping genes were involved in various metabolic processes, cellular transport, and aging. However, one of the most male-biased genes with elevated Fst (0.41) was doublesex and mab-3 related transcription factor 1 (*dmrt1*) – another well-known sex-determination gene in teleosts.

As we sampled blood-rich organs, the expression profile should not solely cover the specialized processes in the organs themselves but also show some of the blood expression patterns. We examined the lists of SBGs with elevated Fst manually for all species and organs (Supplementary tables S12-22). Their putative functions were deduced based on the information on the Zfin database (Bradford et al. 2022) followed by a literature review. It is important to note, that it is difficult to determine their role in a SA relationship due to the interdependence of cellular processes and the pleiotropic nature of genes (Stearns 2010). Furthermore, the roles of SBGs and the reasons for differential expression in a particular organ might not be directly related to the reasons for elevated Fst values at these loci. When discussing the functions of the genes, we thus paid more attention to their potential roles in SA relationships.

In HE, among the SBGs with the highest log fold-change and Fst in both organs was *amh* – one of the key genes involved in vertebrate gonadal differentiation pathway (Morrish and Sinclair 2002; Yatsu et al. 2016; Nagahama et al. 2021; Wagner et al. 2023). Besides that, *amh* also appears to have functions in the brain (Pfennig et al. 2015; Silva and Giacobini 2021).

Among the SBGs two genes are involved in spermatogenesis. The DEAD (Asp-Glu-Ala-Asp) box polypeptide 4 (*ddx4*) appeared within SBG/Fst lists in both organs. It is an RNA helicase, mostly known as a germline factor (Raz 2000; Hickford et al. 2011; Hansen and Pelegri 2021), has functions outside of germ cell-related processes(Yajima and Wessel 2011; Lasko 2013; Schwager et al. 2015; Yajima and Wessel 2015; Noyes et al. 2023). The exosome component 10 (*exosc10*) was shown to be important for spermatogenesis. Mutant mice with inactivated *exosc10* in male germ cells had small testes, showed impaired germ cell differentiation and were subfertile, while in female mice *exosc10* was found to be important in oogenesis (Jamin et al. 2017; Demini et al. 2023). The *exosc10* is also involved in inactivation of X chromosome in somatic cells via regulation of Xist expression (Ciaudo et al. 2006).□

Several genes under ongoing SA were involved in synaptic and neuronal processes. The low-density lipoprotein receptor-related protein 4 (*lrp4*) is one of the regulators of the Wnt/β-catenin signaling pathway and is important in the vertebrate neuromuscular junction (Shen et al. 2015; Mosca et al. 2017; Walker et al. 2021). The gene has other roles, namely in bone homeostasis as shown in mice (Chang et al. 2014; Shen et al. 2015), and roles in tooth development as shown in zebrafish (Ahn et al. 2017). The interleukin 1 receptor accessory protein-like 1b (*il1rapl1b*) belongs to a family of IL1/Toll receptors, regulates dendrite and synapse formation and is responsible for the X-linked intellectual disabilities in humans (Yamagata et al. 2015; Montani et al. 2018). *Il1rapl1*-knockout mice showed mildly impaired working memory and remote fear memory, high locomotor activity, and decreased open-space and height anxiety (Yasumura et al. 2014). In zebrafish *il1rapl1b* is involved in synapse formation in olfactory sensory neurons (Yoshida and Mishina 2008). Another gene on the list involved in regulation of the Wnt/β-catenin signaling – Van Gogh-like (Vangl) planar cell polarity protein 1 (*vangl1*) (Mentink et al. 2018). Certain variants of vangl1 are associated with neural tube defects in zebrafish, humans and mice (Torban et al. 2008; Kibar et al. 2009; Reynolds et al. 2010; Yang et al. 2017). The Triple-A syndrome (*aaas*) gene was among the genes under ongoing SA in HE. Mutations in *aaas* are responsible for the Allgrove syndrome in humans which is associated with multiple neurological abnormalities (Handschug 2001; Brooks et al. 2004; Bizzarri et al. 2013).

Two immune-related genes also appeared among genes under ongoing SA, namely protein tyrosine phosphatase non-receptor type 22 (*ptpn22*) – negative regulator of T cell receptor signaling variants which are associated with severe autoimmune diseases (Fousteri et al. 2013; Armitage et al. 2021; Tizaoui et al. 2021), and limb development membrane protein 1-like (*lmbr1l*) – a gene that was shown to be essential for lymphopoiesis in mice, and mutations in which causing severe immunodeficiency (Choi et al. 2019). Interestingly, *lmbr1l* modulates lymphopoiesis via negative regulation of the Wnt/β-catenin signaling pathway. □

In NO, there were several genes under ongoing SA involved in DNA break repair, and chromosome maintenance, inactivation, and DNA replication. The tonsoku-like (*tonsl*) gene is a DNA repair protein, certain variants of which cause abnormalities in skeletal development (Burrage et al. 2019). Moreover, *tonsl* was found to be regulating epithelial cell immortalisation in breast cancer (Khatpe et al. 2023). The minichromosome maintenance complex component 4 (*mcm4*) as part of the minichromosome maintenance (MCM) complex is responsible for DNA replication (Sheu et al. 2014). In humans, a mutation in *mcm4* causes genomic instability and natural killer cell deficiency (Gineau et al. 2012). The partner and localizer of BRCA2 (*palb2*) – is an important DNA double-strand break gene in humans, mutations in *palb2* are associated with breast cancers (Wu et al. 2020). In mice *palb2* male mutants showed reduced fertility(Simhadri et al. 2014), and in zebrafish *palb2* (*fancn* in the publication) knockout caused female-to-male sex reversal phenotype (Ramanagoudr-Bhojappa et al. 2018).

The H2A.X variant histone (*h2ax*) – a histone variant that is one of the key actors in DNA break repair and chromatin remodeling (Pinto and Flaus 2010). The *h2ax* is involved in sex chromosome inactivation during meiosis in male mice (Fernandez-Capetillo et al. 2003; Turner et al. 2004).

The leucine-rich repeats and WD repeat domain containing 1 (*lrwd1*) is presumed to be the part of origin recognition complex (ORC). ORC is responsible for the initiation of DNA replication and DNA repair (Shen et al. 2010; Vermeulen et al. 2010; Liu et al. 2023).

The final gene in this group, RAD51 paralog D (*rad51d*), is a member of the RAD51 recombinases that are highly conserved across species (Lin et al. 2006; Greenhough et al. 2023). For successful recombination RAD51 is required to interact with *palb2*, which assists localization of it to DNA damage sites (Zhang et al. 2009; Greenhough et al. 2023). Mutations in *rad51d* are associated with a high risk of ovarian cancer in humans (Suszynska et al. 2020; Greenhough et al. 2023). Besides DNA damage repair and replication, one of the genes – ATPase sarcoplasmic/endoplasmic reticulum Ca2+ transporting 3 (*atp2a3*) was found to be among genes under ongoing SA related to hypoosmotic regulation in Takifugu species (Zhang et al. 2020).

#### 3.4.4 Intersex SNP densities and kmerGWAS results in *Hippocampus erectus*

Our analyses identified HE as the most prominent candidate for a sex chromosome SD system among the investigated species. We identified three scaffolds as putative sex chromosomes: *chr14*, *chr18*, and *chr1*. The *chr14* has the highest numbers of SSAs and DEAs as well as the highest number of elevated Fst windows, which makes it the most likely candidate. At the same time, the classical sex-determination gene – *amh*, was among the genes with the highest Fst values and showed the highest expression bias in males. The *amh* is located on scaffold *chr13* with an overall low Fst.

In order to give additional support to *chr14* as HE sex chromosome, we compared male/female SNP densities across the genome and ran a kmerGWAS analysis (the SexFindR workflow) to exclude that scans on unphased genomes might have introduced bias in the results, since the latter does not rely on genome information. Both analyses provided similar results, highlighting *chr14* as a definite sex-chromosome candidate. The results from male/female SNP density scans were much noisier. The KmerGWAS analysis resulted in a clear peak for *chr14* (Figure 3B).

**Figure 3.**
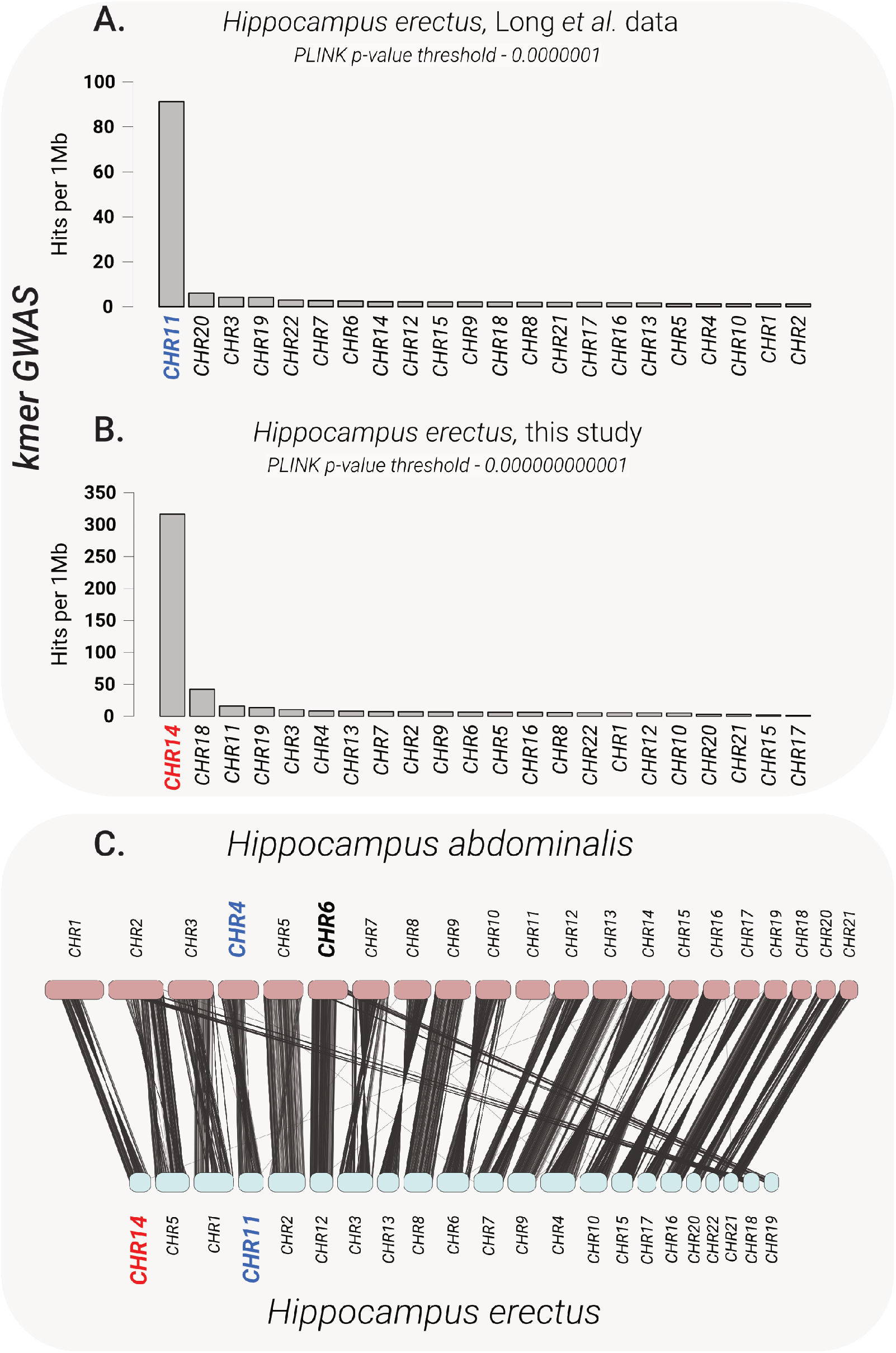
Panels A and B - results of kmerGWAS analysis. Y-axis - number of ABYSS contigs that aligned to the genome with BLASTn per 1Mb (i.e. adjusted for size), X-axis - the chromosome. **Panel C** - synteny analysis between *H.abdominalis* and *H.erectus* genome. Each line connects 1-1 orthologues in HA and HE. HA chromosomes were ordered based on their length, HE chromosomes were ordered by their relation to HA genome. Red marks sex-linked chr identified in our study. Blue - HE sex-linke chr identified by Long, Bold Black - *H.abdominalis* sex-linked chr.

#### 3.4.5 Long et al. data comparison

Long and colleagues had recently published a study investigating sex chromosomes in three syngnathids, including HE. In that study, the HE genome was reconstructed using the genome of *Hippocampus abdominalis* (HA), a phylogenetically quite derived species compared to HE (Betancur-R et al. 2017) with a larger genome that underwent an additional chromosome fusion (He et al. 2022; Long et al. 2023).

To compare our results to Long et al. 2023, we performed a synteny analysis between the HA genome used by Long and the HE genome at our disposal (PRJNA613176) (Li et al. 2021). For this purpose, we used annotation data from these genomes, identified 1–1 orthologs, and used the locations of these orthologs in each genome to identify homologous chromosomes (Figure 3C). The sex chromosome that we identified (*chr14*) and *H. abdominalis chr4* identified as sex chromosome in HE by Long et al. were not homologous. For additional tests, the data published by Long et al. 2023 was used. Here we again opted for kmerGWAS because it gave us the cleanest signal, and because identification of sex-linked kmers is not affected by assembly quality. The resulting sex-associated kmers were aligned to the HE genome. The results of kmerGWAS confirmed Long’s finding that *chr4* (HA)/*chr11* (HE) is sex-linked, although the signal was not as pronounced as in HE using our data (Figure 3A). This suggests that in HE, just like in some other teleost species, sex chromosomes are non-stable and could be population/strain dependent.

## 4. Discussion

We evaluated the signatures of active sexual antagonism (SA) as well as the putative mechanism of sex determination (SD) in three syngnathid species that evolved on a gradient of male pregnancy and sex-role reversal. Our objective was to determine whether the pronounced SA pressure, caused by male pregnancy and shifts in sex roles, can directly lead to the emergence of distinct sex-linked genomic regions. Alternatively, SA and SD would have no causal relationship between each other but evolve in parallel.

### 4.1 Sexual antagonism in three syngnathid species along the male pregnancy gradient

Based on prior SA research in other species we hypothesized that there would be a subset of genes experiencing unresolved intralocus sexual conflict, meaning that the divergent fitness effects of these genes in males and females could not be mitigated solely through differential gene expression or genetic fixation. We focused on two main indicators of SA: I) Intersex expression bias – differences in gene or allele expression between males and females that suggest intralocus sexual conflict resolution via sex-specific optimal expression levels. II) Intersex genetic divergence – genetic divergence measured by the fixation index (Fst) signifies resolution of intralocus conflict via confinement of SA alleles to the benefiting sex. Loci that exhibit both indicators should be considered markers of ongoing SA and are key to understanding the roles of SA in genetic divergence between sexes and the underlying biological processes that drive the SA relationships.

As an alternative signature of ongoing SA, our analysis included investigation of differentially expressed alleles (DEAs) and sex-specific allele (SSA) expression. These mechanisms often remain unnoticed in SA and SD studies, which primarily concentrate on biased gene expression and genetic divergence signatures alone. Instead of restricting conflicting alleles to one sex through recombination suppression, each sex could express only the beneficial allele, eliminating the need for allele fixation.

Male pregnancy and sex-role reversal likely amplify the divergence in fitness strategies between the sexes, thus intensifying SA pressures. We propose that this SA-induced genetic divergence may influence the evolution of sex determination (SD) systems in syngnathids.

#### 4.1.1 Genes under ongoing SA. SBG-Fst overlap

In our study, we investigated three syngnathid species with contrasting brooding strategies that exist along the male pregnancy gradient: *Nerophis ophidion* (NO) with basic external egg attachment, *Syngnathus typhle* (ST) with an inverted brood pouch and a placenta-like structure, and *Hippocampus erectus* (HE) with a sealed brood pouch and a placenta-like structure.

The degree of male pregnancy was hypothesized be the key driver of the SA relationship and consequently affect the evolution of SD systems within these species. We expected NO to show the weakest SA signal. ST to show intermediate SA signal, with a moderate number of active SA markers and possibly a small sex-linked region or loci. And HE, with the most advanced pregnancy form, was predicted to show the strongest SA signals, including a large sex-linked genomic region rich in pregnancy-related genes.

Alternatively, we considered that sex roles and mate competition would be major drivers of SA and SD system evolution. In our study, HE represents a species with traditional sex roles. We expected a higher count of active SA markers with male bias and signs of an XY SD system, due to HE males experiencing stronger selection. ST and NO represent species with reversed sex roles, with females under stronger selective pressure. Thus, we expected to see more SA markers with a female bias in these species, and potentially a ZW SD system. This picture is complicated when we factor in mating strategies (monogamous or polygamous). Of course, we only considered phenotypes that are known to us, but there could be dimorphism and conflict areas that we haven’t considered, suggesting there might be many alternative patterns.

Furthermore, whether SA and SD formation processes are linked also remains to be determined. It is logical to assume that if SA pressure is strong enough and could not be resolved via bias in expression, it would lead to the formation of a sex-linked genomic region. However, if SA could be primarily resolved via bias in gene or allele expression, then SA and SD system formation processes would probably be unlinked. The data, however, suggests a more nuanced reality than our initial two hypotheses proposed.

The least divergence between the sexes was observed in ST, as indicated by both the overall intersex Fst levels across the genome and the numbers of SBGs with elevated intersex Fst values. This hints that the resolution of SA conflict primarily occurs through sex-specific gene and/or allele expression, or through other unknown mechanisms. An elevated Fst of 0.4 for *dmrt1*, a known SD gene (Kobayashi et al. 2013), suggests ongoing intralocus sexual conflict for this SBG. Despite other genes exhibiting higher Fst values in different genomic locations and the nearly equal numbers of SSAs between the sexes, *dmrt1* remained a robust SD gene candidate for ST. This is not only because SD predominantly involves co-option of the same set of genes, but also as its Fst value is close to the expected 0.5 for a fixed difference between sexes (Gammerdinger et al. 2020).

In HE, SBGs with elevated Fst clustered on a single chromosome-length scaffold, suggesting a putative sex chromosome. Notably, several genes associated with synaptic and neuronal processes were found among these SBGs. This reminds of a recent study on signatures of SA on the sex chromosomes of *Gasterosteus nipponicus* (Japan Sea stickleback) that identified genes associated to mental disorders and neurological human development near regions of high intersex Fst on the sex chromosome (Dagilis et al. 2022). The role of these genes in SA, however, remains as of yet undiscovered.

In NO, the SBGs with elevated Fst were scattered throughout the genome and did not cluster within a particular genomic region. Overall, the genome itself showed elevated intersex Fst levels suggesting reduced recombination landscape in this species or sex specific lethality. The notable groups of genes showing signatures of ongoing SA include genes associated with DNA break repair, chromosome maintenance, inactivation, and DNA replication. While it is tempting to speculate that their involvement in SA could be linked to spermatogenesis, given the importance of these genes in many cellular processes, it remains difficult to identify their roles in SA relationships in NO.

Our analysis of SBGs with elevated intersex Fst unveils that species with traditional sex roles and the most advanced form of male pregnancy (HE) exhibit the strongest SA signature and male bias which is consistent with both our hypotheses. In contrast, ST with its intermediate male pregnancy and sex-role reversal, shows the weakest SA signal, while NO, featuring the basic form of male pregnancy and extreme sex-role reversal, presents a moderate SA signal alongside notable genetic divergence between sexes. While mate choice is somewhat biased towards the male sex, i.e. males are the choosing sex, in both ST and NO, e.g., (Berglund and Rosenqvist 1993), females in ST have a more pronounced role in mate choice compared to NO females. This proposes sexual selection on females in ST to be less skewed than in NO, potentially allowing it to deal with SA loci via expression bias alone. It has been suggested that differences in mate choice and secondary sexual characteristics between NO and ST may reflect their distinct egg production regimes (Sogabe and Ahnesjö 2011). In NO, the eggs are produced in cycles, therefore mating system and secondary sexual characteristics are tailored to signal immediate availability for mating (Sogabe and Ahnesjö 2011). On the other hand, in ST the egg production is asynchronous, meaning that most ST females would be able to deposit some eggs whenever potential mates are available (Sogabe and Ahnesjö 2011). This mating strategy should impose selection on traits and pathways that enable the ongoing opportunity for spawning.

Our findings suggest that neither male pregnancy, nor sex roles are the sole drivers of SA across these species. Rather, the dynamics of sex roles and mate choice are pivotal, with the sex subjected to higher competition being under stronger selection and showing higher signatures of active SA. Notably, the elevated genetic divergence that we observed in NO with no distinct sex-linked region suggests alternative, to ST and HE, mechanisms that mediate SA selection.

#### 4.1.2 Non-coding RNAs as mediators of sexual antagonism in syngnathids. Evidence from DEAs

In all three syngnathid species, non-coding RNA processing and metabolism were among the pathways that exhibited differential allele expression between the sexes. Enrichment of the GO term “ncRNA metabolic process”,identified in all examined species, suggests that genes involved in chemical reactions and pathways related to non-coding RNA transcripts are under ongoing SA pressure and require sex-specific allele expression optima (Ashburner et al. 2000; The Gene Ontology Consortium et al. 2023). We observed a similar enrichment pattern in ncRNA-related pathways when we looked for intersex differences in gene expression in old and young ST individuals (Pappert et al. 2023). Taken together, this suggests an active role of ncRNAs in SA processes in syngnathids. The function of ncRNAs in SA processes could be especially important in species with low genetic divergence between the sexes like in ST.

Non-coding RNAs that are involved in translation processes include ribosomal RNAs (rRNAs) and transfer RNAs (tRNAs). The regulatory ncRNAs are associated with various cellular processes and include: long-noncoding RNAs (lncRNAs), microRNAs (miRNAs), piwi-interacting RNAs (piRNAs), small nuclear RNAs (snRNAs) and small nucleolar RNAs (snoRNAs) (Frías-Lasserre and Villagra 2017; Bartel 2018; Kopp and Mendell 2018; Mattick et al. 2023). A plethora of reports discuss the effects of regulatory ncRNA expression between the sexes on development, physiology, immunity and disease (Zhou et al. 2014; Gershoni and Pietrokovski 2017; Zhang et al. 2017; Gal-Oz et al. 2019; Rosspopoff et al. 2023; Wang et al. 2023).

Noncoding RNAs have an important role in pregnancy progression. miRNAs regulate the physiological processes in a sexually dimorphic manner, ensuring normal fetal development, successful pregnancy, and susceptibility to disease (Varì et al. 2021). A known role of ncRNA roles in sexual development is the inactivation of the with XIST in humans and other placental mammals (Brown et al. 1991; Lee et al. 1999; Dupont and Gribnau 2013; Posynick and Brown 2019; Rosspopoff et al. 2023). However, much of the literature on sexually antagonistic co-evolution and intralocus sexual conflict in non-model organisms focuses on either fixed genetic differences or differential gene expression as a way to resolve intralocus sexual conflict, while the roles of ncRNAs in SA relationships remain as yet understudied. Epigenetic mechanisms are also responsible for governing gene-hormonal-environmental interactions. There is a considerable amount of data from teleost fish showing that miRNAs and lncRNAs are involved in sex determination processes and gonadal differentiation (Bizuayehu et al. 2012; Juanchich et al. 2013; Jing et al. 2014; Lau et al. 2014; Xiao et al. 2014; Juanchich et al. 2016; Tao et al. 2016; Wang et al. 2017; Pinhal et al. 2018; Qiu et al. 2018; Yan et al. 2021). Studies have also brought up the potential roles of ncRNAs in the regulation of immune responses in teleost fish (Wang et al. 2018).

In addition to sex-specific effects via regulatory ncRNA, the potential roles of ncRNAs in mediating SA could be related to expression and processing of rRNA and tRNA. The 5S and 18S rRNA abundance, and 5S–18S rRNA ratio have been used as sex markers in fish (Shen et al. 2017). Though Shen et al. discuss development of ovaries and testes, variation in rRNA transcript abundance can have differential importance for sexes in other tissues. Differential rRNA allele expression could be another mechanism mediating SA relationships in syngnathids. Tissue-specific rRNA allele expression has been described in humans and mice (Parks et al. 2018). Transcending differential transcript abundance and allele expression, RNA’s ability to form specific structures is hypothesized to have an impact on epigenetic differences between the sexes as exemplified by a study on variation in R-loop formation in *Drosophila melanogaster* (Stanek et al. 2022).

#### 4.1.3 Highly divergent alleles may reflect allele-specific expression, not fixation

Our Weir Fst analysis, based on 10kb windows, is expected to show Fst values around 0.5 for completely fixed differences between homozygous and heterozygous sexes (Gammerdinger et al. 2020). While we expected a slightly lower mean Fst due to windowing, surprisingly, genes like *irf2* and *amh* (HE), *plp1b*, *ttc3*, *tcnbb* in (NO), and *trak2* (ST) displayed Fst values exceeding 0.8, suggesting near-complete fixation of opposite alleles in each sex. A more likely explanation for the observed extreme Fst values is either highly biased or allele-specific expression (ASE). ASE can arise from chromosome inactivation, regulatory elements, nonsense-mediated decay, or imprinting (Castel et al. 2015; Fan et al. 2020). While extensively studied in human pathology (Langmyhr et al. 2021; Rao et al. 2021; Van Beek et al. 2023), sex-specific ASE in the context of SA selection remains underexplored.

### 4.2 Unraveling sex determination systems in three syngnathids

This study aimed to identify SD systems in three syngnathid fish species (NO, ST, and HE) and investigate their relationship to loci under ongoing SA pressure. In HE, we observed an over-accumulation of SBGs with and without elevated Fst, DEAs, and SSAs on a single chromosomal scaffold, suggesting a putative sex chromosome. However, in the other two species (NO and ST) our investigation of loci under SA did not show any accumulation within single or multiple distinct genomic regions.

#### 4.2.1 No GSD in pipefishes Nerophis ophidion and Syngnathus typhle?

Genetic sex determination (GSD) systems typically involve either a chromosomal region or entire chromosome linked to one sex, often containing master SD gene/genes. Sometimes, an SD locus can be as small as a single nucleotide polymorphism, like in *Takifugu rubripes* (Kamiya et al. 2012). Regardless of the size of the sex-linked region, their identification usually involves scans for intersex Fst, calculation of differential coverage statistics, search for SBG overaccumulation, GWAS for sex-linked alleles, or all of the above.

Despite relatively high genome-wide intersex Fst in NO, distinct regions with elevated Fst and/or regions with accumulation of SBGs were absent. Notably, several genes exhibited extreme Fst values but they were scattered throughout the genome. None these genes had known SD functions, which suggests that allele-specific expression may be responsible for extreme Fst values at these loci. NO also possessed both male and female SSAs. All this hints towards an Environmental Sex Determination (ESD) system in NO.

ST showed the lowest genetic differentiation between the sexes and no distinct regions of elevated divergence. Also, ST showed the lowest number of SBGs, and the lowest number of DEAs among studied species. These patterns, reinforced further by similar findings in *Syngnathus scovelli* (Long et al. 2023), could hint towards an ESD system. However, the presence of a well-known teleost SD gene (*dmrt1)*, with an Fst of 0.4 complicates the picture. The Fst of 0.4 suggests that one sex is homozygous while the other is heterozygous at this locus, pointing towards an XY/ZW sex determination system with *dmrt1* as the master SD gene. Further investigation is needed to determine SD mechanism in ST.

interestingly, recent research on *Ictalurus punctatus* (channel catfish) (Bao et al. 2019) revealed a novel mechanism where sex is determined via expression of a male-specific isoform of breast cancer anti-resistance 1 (*BCAR1*) gene during early developmental stages. Just like sex-specific isoform expression, DEA and ASE may represent hidden SD mechanisms in species with putative ESD systems, such as NO or ST.

#### 4.2.2 Intraspecies sex chromosome polymorphism in HE and multi-chromosome SA supergene

Our study explored signatures of ongoing SA and their relationship to the SD systems in three syngnathid species. In HE, our analysis revealed that a single chromosome-level scaffold (*chr14*) accumulated the majority of ongoing SA indicators: SBGs with elevated Fst, DEAs, and SSAs. This scaffold also showed an overall elevated Fst levels (0.5 or above), with males being a heterozygous sex, which suggest an XY SD system. These patterns hint that in HE, ongoing SA relationships might be the driving force behind an increased genetic divergence between the sexes, which lead to the formation of a sex-linked chromosome.

A sex-linked chromosome was recently described in HE (Long et al. 2023). The sex-linked chromosome that we identified, however, was not homologous to the homomorphic sex-linked chromosomes described in *Hippocampus erectus* (chr11 or homologue of *Hippocampus abdominalis* chr4) and *Hippocampus abdominalis* (chr6) (HA) with low sex-specific sequence divergence suggesting a recent evolutionary trajectory (Long et al. 2023). The identified discrepancies in HE sex chromosomes may have either appeared due to differences in methodological approach, or alternatively indicate real biological differences of the two HE breeds.

Methodological differences in the two studies become evident in several aspects: i) the type of sequence data utilized – this study employed RNA data for identifying the sex-linked chromosome, whereas Long et al. used DNA; ii) the genome used for mapping the generated sequencing data – this study used the chromosome-level HE genome, whereas Long et al. used the genome of the more distant HA; and iii) the tools and metrics utilized for identifying sex-linkage – we focused on over-accumulation of SA signature on particular genomic region, while the other study relied on Fst and GWAS analyses. The bioinformatic tools that were used to conduct the analysis were different, which could contribute to differing results.

To enhance the comparability of the two datasets and potentially mitigate the impact of methodological differences as explanations for the disparities in HE sex chromosome identification, we re-analyzed the data from HE males and females as presented in Long et al., employing the chromosome-level assembly of HE from (Li et al. 2021) with kmerGWAS. Using this approach, we have discovered compelling evidence supporting the evolution of two distinct sex-linked chromosome systems in the two breeds of HE, facilitating the resolution of SA. Previously, differences in sex-linked regions between captive breeds and wild-types have been observed in various species, such as zebrafish, Siamese fighting fish and cichlids (Wilson et al. 2014; Panthum et al. 2022; Wang et al. 2022; Lichilín et al. 2023; Schartl et al. 2023). For instance, the Z chromosome of the zebrafish wild-type ZW SD system was hypothesized to have been lost during domestication (Wilson et al. 2014; Schartl et al. 2023). The domesticated zebrafish WW individuals develop either as females or neomales, however, their SD mechanism remains as yet unknown (Schartl et al. 2023). Additionally, in Siamese fighting fish, some breeds possess a monogenic SD system with *dmrt1* as a master SD gene, while others exhibit polygenic SD mechanisms (Panthum et al. 2022; Wang et al. 2022; Schartl et al. 2023).

The evolution of two diverging sex chromosome systems, in two lineages of domesticated HE lineages suggests an even younger evolutionary age of the sex-linked chromosomes than previously proposed by Long et al. (Long et al. 2023). The HEs used in Long et al. originated from a cultured population in Fujian (China), with a seed population of 2,000 – 3,000 wild-caught individuals. In contrast, HEs used in this study were obtained from a German aquarium breeder, with an unknown but likely smaller population size. The emergence of the two sex chromosome systems may have occurred in a similar scenario as reported for captive zebrafish, where domestication selected for different SD loci than in captive populations (Schartl et al. 2023). However, the incomplete linkage in both sex chromosomes, as indicated by the Fst in our case (less than 0.4) and by the GWAS in Long et al. (Long et al. 2023) (male homozygotes were found within a sex-linked region), raises questions concerning the existence of a master SD locus, while the recombination landscape could predate a turnover event as proposed previously (Long et al. 2023).

Genes with sex-biased expression patterns often exhibit elevated intersex Fst values (Cheng and Kirkpatrick 2016; Wright et al. 2018; Lichilín et al. 2021). In humans and flies, this relationship follows a “Twin Peaks” pattern. Genes with intermediate bias in expression demonstrate the highest Fst values, indicating ongoing SA, whereas significant bias in expression aids in resolving SA (Cheng and Kirkpatrick 2016). However, in cichlids and guppies, the intersex Fst increases with expression bias, thus, the most extreme sex-biased genes display the highest intersex Fst values (Lichilín et al. 2021; Wright et al. 2018). These data indicate that a bias in expression alone may not be sufficient to fully resolve sexual conflict, independent of the underlying mechanism.

Notably, despite its young age, we observed an enrichment of SBGs on the HE sex-linked chromosome compared to the rest of the genome. This, combined with an elevated intersex Fst pattern, as revealed by kmerGWAS or GWAS as in Long’s study, suggests the presence of a true sex chromosome. The elevated intersex Fst values likely result from ancestral low recombination rates, and Long et al. proposed several promising SD candidate genes in this context (Long et al. 2023).

In this study we were unable to identify SD candidates related to the canonical vertebrate gonadal differentiation pathway on our sex-linked chromosome. Instead, we observed extreme intersex Fst (0.9) and extreme bias in expression for *amh*, a well-known sex determination gene, which is however not located on the sex chromosome candidate of HE This observation suggests a transient state during a turnover event, as recently suggested for multi-locus SD systems (Schartl et al. 2023).

In a plausible evolutionary scenario, the chromosome containing the *amh* gene as an SD locus was originally the only sex-linked chromosome in HE. The a*mh* falls within the canonical vertebrate gonadal differentiation pathway and genes related to *amh*, like anti-Müllerian hormone type II receptor (Nakamoto et al. 2021), were identified as SD loci in other syngnathid species (Qu et al. 2021). Domestication in closed aquarium systems, devoid of natural predators and subjected to selection distinct from those in the natural population, likely amplified the effects of SA. This led to an enrichment of the now identified sex-linked chromosome for SBGs with elevated Fst values. Although unusual for such a young sex chromosome, selection for elevated Fst values was likely facilitated by ongoing intralocus sexual conflict that caused bias in expression promoting genetic linkage. Fixation must have been accelerated by captive breeding with a limited number of mates resulting in an overall reduced recombination landscape within the genome that facilitated linkage of *chr14* (or *chr11* in the other HE breed). While a new chromosome became sex-linked, the master SD locus (*amh*) remained the same, creating a multi-locus SD system. This could be similar to recently described multi-chromosome supergene in ants that contains socially antagonistic alleles (Scarparo et al. 2023), however, in case of HE it rather contains sexually antagonistic alleles.

Several questions remain unanswered, including the speed of such transformation, the exact mechanisms for recombination slowdown, the relationships between bias in expression and elevated Fst, and the role of ongoing SA in this process. However, if confirmed, populations of *Hippocampus erectus* offer a valuable system to study the effects of SA on genome evolution.

## Conclusion

Overall, we found that sex roles and mate competition dynamics, rather than male pregnancy, are the major drivers of sexual antagonism (SA) in the studied syngnathid species. Our initial hypothesis suggested that a strong SA selection pressure, e.g., exerted by male pregnancy, may be enough to lead to the formation of a genetic sex-determination (GSD) system. We found no such direct link in *Syngnathus typhle* and *Nerophis ophidion*. At the same time, in a species with the most derived form of male pregnancy, *Hippocampus erectus*, we identified a putative sex chromosome which also exhibited over-accumulation of ongoing SA markers. This may indicate, that in *H. erectus* SA relationships have been amplified by the adaptation to male pregnancy. However, the identified sex-linked chromosome was not specifically enriched in pouch-related genes. It seems that, while male pregnancy might have amplified the SA, it was not the major driver behind the sex-linked chromosome’s formation. Notably, we observed what appears to be an intraspecies sex chromosome polymorphism in different breeds of *H. erectus*, which possibly evolved due to prolonged captive breeding. In such captive-bred systems, or natural populations experiencing bottlenecks, SA selection pressure may be especially important in driving genetic divergence between the sexes. In all species we also discovered that processes related to non-coding RNAs and allele-specific expression seemed to be important mediators of SA, marking them as targets for future SA research.

## Ethics statement

Work was carried out in accordance with German animal welfare law and with the ethical approval given by the Ministerium für Energiewende, Landwirtschaft, Umwelt, Natur und Ditgitalisierung (MELUND) Schleswig-Holstein (permit no. V242-57983/2018). No wild endangered species were used in this investigation.

## Authors’ contributions

A.D., O.R. and A.B. planned the project. J.P. collected and processed samples. A.D. analyzed the data and interpreted the results with input from O.R. and A.B.. A.D. & O.R. wrote the manuscript with input from all coauthors.

## Data Access

All sequencing data generated in this project can be found under BioProject ID PRJNA947442.

